# “The Type IV Secretion System of *Patescibacteria* is homologous to the bacterial monoderm conjugation machinery”

**DOI:** 10.1101/2025.01.22.634366

**Authors:** María del Mar Quiñonero-Coronel, Pedro J. Cabello-Yeves, Jose M Haro-Moreno, Francisco Rodríguez-Valera, M. Pilar Garcillán-Barcia

## Abstract

The Candidate Phyla Radiation, also known as *Patescibacteria*, represents a vast and diverse division of bacteria that has come to light via culture-independent “omics” technologies. Their limited biosynthetic capacity, along with evidence of their growth as obligate epibionts on other bacteria, suggests a broad reliance on host organisms for their survival. Nevertheless, our understanding of the molecular mechanisms governing their metabolism and lifestyle remains limited. The Type IV Secretion System (T4SS) represents a superfamily of translocation systems with a wide range of functional roles. T4SS genes have been identified in the CPR group *Saccharibacteria* as essential for their epibiotic growth. In this study, we used a comprehensive bioinformatics approach to investigate the diversity and distribution of T4SS within the *Patescibacteria* lineage. The phylogenetic analysis of the T4SS signature protein VirB4 suggests that most of these proteins cluster into a distinct monophyletic group with a shared ancestry to the MPF_FATA_ class of T4SS. This class is found in the conjugative elements of *Firmicutes*, *Actinobacteria*, *Tenericutes*, and *Archaea*, indicating a possible horizontal gene transfer from these monoderm microorganisms to CPR bacteria. We identified additional T4SS components near *virB4*, particularly those associated with the MPF_FATA_ class, as well as homologs of other T4SS classes, such as VirB2-like pilins, and observed their varied arrangements across different CPR phyla. The absence of a relaxase in most of these T4SS clusters suggests that the system has been co-opted for other functions in CPR bacteria. The proximity of T4SS components to the origin of replication (gene *dnaA*) in some CPR suggests a potential mechanism for increased expression. The broad ubiquity of a phylogenetically distinct T4SS in CPR, combined with its chromosomal location, underscores the significance of T4SS in the biology of *Patescibacteria*.

**Impact statement:** The Candidate Phyla Radiation (CPR), or Patescibacteria, represents a highly diverse bacterial group that constitutes a significant fraction of the microbial dark matter. Known for their minimal biosynthetic capabilities and dependence on host bacteria, CPR bacteria exemplify fascinating yet poorly understood lifestyles. Our research sheds light on the molecular strategies underlying their survival by focusing on the Type IV Secretion System (T4SS), a versatile bacterial machinery typically involved in DNA transfer and effector molecule delivery. We revealed that T4SS is not only widespread in CPR but also forms a distinct evolutionary lineage, likely acquired via horizontal gene transfer from Gram-positive bacteria, and we illustrated its gene organization across different CPR phyla. Unlike canonical systems, CPR T4SS lacks key components for conjugation, hinting at a novel functional adaptation. Remarkably, T4SS genes in CPR are often located near the replication origin, suggesting a regulatory role linked to their expression. This research suggests the repurposing of T4SS in CPR to thrive under extreme reliance on other organisms. By unraveling the genomic and evolutionary peculiarities of T4SS in CPR, our work provides valuable insights into their unique biology, broadens our understanding of microbial diversity and innovation, and highlights this system as a strong candidate to sustain their parasitic episymbiotic lifestyle.

**Data summary:** The authors confirm all supporting data, code and protocols have been provided within the article or through supplementary data files.

## Introduction

Type IV secretion systems (T4SSs) are multiprotein nanomachines present in both Gram-negative and Gram-positive bacteria. They span the entire bacterial cell envelope and function as a one-step mating-pair formation (MPF) system, capable of delivering effector molecules directly to the cytosol of a target cell, typically requiring direct cell-to-cell contact (*1*). T4SSs are mainly involved in bacterial conjugation and effector secretion from bacteria to eukaryotic cells (*2*). Bacterial conjugation is one of the most prominent mechanisms of horizontal gene transfer. It involves the transport of a nucleoprotein effector from donor to recipient bacteria (*3*). Meanwhile, T4SSs devoted to protein delivery are mainly deployed by intracellular bacterial pathogens to hijack eukaryotic host cell processes, gain access to nutrients, and regulate specific virulence programs to ensure their survival into the host (*4*). While T4SSs known to be involved specifically in protein delivery have been mostly found in diderms, only a few reports show the presence of T4SS protein translocators in monoderms, more specifically in streptococci (*5–7*). More recent studies, however, highlight a broader functional diversification of the T4SS. T4SS-mediated toxin export to other bacteria pointed to an additional role of T4SS in killing other bacteria (*8–11*). Moreover, there are a few examples of T4SSs capable of importing or exporting DNA from or to the environment (*12–14*). Furthermore, some reduced T4SS variants have been described in *Archaea,* the Ced system in *Sulfolobales* and the Ted system in *Thermoproteales*, mediating the unidirectional import of DNA between archaeal cells, which is then used as a template for genome repair by homologous recombination (*15–17*).

Understanding the role of T4SS in different bacterial lineages is crucial for uncovering main aspects of their physiology and cellular biology. However, in other bacterial groups outside of *Proteobacteria* and *Firmicutes*, T4SSs are poorly characterized. One of these underexplored groups is *Patescibacteria*, also known as Candidate Phyla Radiation (CPR), which encompasses a strikingly diverse collection of ultra-small bacteria present in a variety of habitats with streamlined genomes (*18*). This bacterial division was placed as a member of the *Terrabacteria* clade and a sister lineage to the *Chloroflexota* and *Dormibacterota* branches, suggesting that CPR evolved by reductive genome evolution from free-living ancestors (*19*). CPR members are typically characterized by extremely limited biosynthetic capacities, including the inability to synthesize amino acids, lipids, vitamins, and, in many cases, nucleotides (*20–22*). Additionally, they lack genes for the citric acid cycle and oxidative phosphorylation, except for the ATPase complex. As a result, a symbiotic dependence for cellular growth has been assumed, although free-living and particle-associated groups have been detected (*23*). Most knowledge on CPR has been obtained through DNA-based, culture-independent methods because they are generally recalcitrant to traditional cultivation techniques. However, a few strains, primarily from the CPR phylum *Saccharibacteria* (*24–30*), as well as few from *Gracilibacteria* (*31, 32*), *Absconditabacteria* (*33*), and *Paceibacteria* (*34*), have been successfully cocultured with a compatible bacterial or archaeal host. Additionally, CPR growing attached to host bacterial surfaces in an epibiotic association has been reported, such as the parasitic relationship between *Saccharibacteria* species and various *Actinobacteria* (*24*, *30*), as well as between *Candidatus Vampirococcus lugosii*, a member of the CPR phylum *Absconditabacteria*, and an anoxygenic photosynthetic gammaproteobacterium (*32*).

The presence of a putative T4SS in CPR was first reported in the parasitic *Saccharibacteria Ca. Nanosynbacter lyticus* strain TM7x (*35*) and later in another *Saccharibacteria* species, *Ca. Southlakia epibionticum* strain ML1 (*36*). In this latter strain, a transposon-insertion sequencing genome-wide screen revealed that the T4SS cluster was essential for its epibiotic growth (*36*). These T4SSs have also been identified in CPR metagenome-assembled genomes (MAGs) (*37*) and complete genomes from Lake Baikal (*38*), the largest and deepest freshwater lake on Earth. This study was particularly motivated by the enigmatic and still poorly understood biology and ecological significance of this lineage, which colonizes most habitats on Earth. Here, we conducted a lineage-wide analysis of the presence and diversity of T4SS in *Patescibacteria* spanning different habitats and identified the main peculiarities derived from this secretion system.

## Materials and Methods

### Dataset

Assemblies from the *Patescibacteria* group (4,699) were downloaded from NCBI RefSeq database (version 212, March 2023) and only those confirmed as members of *Patescibacteria* in the Genome Taxonomy Database (GTDB, release 214) (*39*) (3,024 assemblies) were kept. Additionally, *Saccharibacteria Ca. Southlakia epibionticum* strain ML1 (NZ_CP124550.1) and *Ca. Nanosynbacter lyticus* strain TM7x (NZ_CP007496.1), for which the presence of a T4SS was reported (*35*, *36*) were included. Furthermore, nine complete CPR genomes isolated from the Baikal Sea (*38*) were also included (PRJNA924152). The final CPR dataset contained 3,035 *Patescibacteria* genomes, of which 53 complete genomes (Supplementary Table S1 and Supplementary Figure S1).

### Detection of T4SS components

Hidden Markov Model (HMM) profiles of the protein components of the eight phylogenetic classes of T4SS, as described by (*40*) and available at https://github.com/macsy-models/CONJScan/tree/main/profiles, as well as the Pfam HMM profiles TrbC/VirB2 (PF04956.16) and T4SS_pilin (PF18895.3), were used to inspect the *Patescibacteria* protein dataset. Searches were conducted using the *hmmscan* function of HMMER 3.1b2 (*41*), with i-e-value < 0.001 and HMM profile alignment coverage > 50% as inclusion criteria. The presence of relaxases associated with T4SS was checked using MOBscan (*42*).

### VirB4 phylogeny

The VirB4 homologs recovered from the CPR dataset were used for the phylogenetic reconstruction. Additionally, 262 VirB4-like proteins from bacterial conjugative plasmids and integrative and conjugative elements (ICEs) encoding a T4SS of the MPF_FATA_ (227), MPF_FA_ (5 proteins), MPF_T_ (5), MPF_F_ (5), MPF_G_ (5), MPF_C_ (5), and MPF_B_ (5) classes were included (Supplementary Table S4). Furthermore, 12 VirB4-like proteins from *Archaea* were included (5 from the Ced system, 4 from the Ted system, and 3 from conjugative archaeal plasmids belonging to MPF_FATA_) (Supplementary Table S2). Proteins were aligned using MAFFT v7.310 with the L-INS-i method (*43*), and the alignment was curated with Trimal 1.2rev59 (option -automated1) (*44*). A maximum-likelihood phylogenetic reconstruction was generated using IQ-TREE version 1.6.1 (*45*), based on the substitution model LG+F+R10, which was selected as the best fit according to the Bayesian Information Criterion (BIC) provided by ModelFinder (*46*), Branch support was estimated with ultrafast bootstrap (UFBoot) approximation (*47*) and SH-like approximate likelihood ratio test (SH-aLRT) (*48*), using 1,000 replicates in both cases. The resulting tree was midpoint rooted. It was visualized and annotated using the online tool iTOL (*49*). We computed the patristic distance matrix from the tree using the cophenetic.phylo function in the *ape* package (v5.3) for R (*50*).

### Analysis of the *virB4* gene neighborhood

Coding sequences surrounding the *virB4* gene (20 CDSs upstream and 20 downstream) in the CPR assemblies were extracted whenever a second MPF component was identified. Entrez Protein Family Models, including 12,592 NCBIFAM and 4,486 TIGRFAM profiles (available at https://ftp.ncbi.nlm.nih.gov/hmm/16.0/ (*51*)), as well as 19,632 protein family models of Pfam-A (available at http://ftp.ebi.ac.uk/pub/databases/Pfam/releases/Pfam35.0/), were used to query the CPR proteins with the HMMER 3.1b2 *hmmscan* function (parameters -E 0.001 --domE 0.001 -- incE 0.001 –incdomE 0.001) (*41*)). Proteins were also clustered with the *mmseqs* cluster module included in MMseqs2 suite (version 13.45111, (*52*)), with a minimum sequence identity of 30% and 70% coverage (parameters --cov-mode 0 --cluster-mode 0 –cluster-reassign). For each cluster of more than 5 members, a multiple alignment was built was MAFFT v7.310 with default parameters (*43*), and trimmed with Trimal 1.2rev59 (option -automated1) (*44*)). These multiple sequence alignments were used to construct an HMM profile for each cluster, with the function *hmmbuild* of HMMER 3.1b2 (*41*). These HMM profiles were compared with the MPF_FATA_ HMM profiles using the *HHsearch* function of HH-suite v3 (*53*). Clincker (*54*) was used to visualize to the *virB4* gene vicinity of representative genomes.

## Results and Discussion

### The phylogenetic distribution of VirB4 in *Patescibacteria*

VirB4 is the only protein with detectable homologs present in all known T4SS and is, therefore, a hallmark of T4SS presence (*55*, *56*). It functions as an ATPase involved in the assembly of the translocation machinery and pilus biogenesis (*2*). Its phylogenetic distribution is the basis for the eight classes of T4SS described in bacteria, namely MPF_T_, MPF_F_, MPF_I_, MPF_G_, MPF_C_, MPF_B_, MPF_FA_, and MPF_FATA_ (*40*, *55*). The first six are distributed among Gram-negative bacteria, while the last two are exclusive to monoderms, with the final one also present in the conjugative systems of *Archaea*. VirB4 was previously reported to be enriched in CPR compared to non-CPR bacteria (*57*). We searched a CPR dataset containing 3,035 *Patescibacteria* genomes, of which 53 complete genomes (Supplementary Table S1 and Supplementary Figure S1), using a VirB4 HMM profile. We retrieved 2,474 homologs in 2,417 out of 3,035 CPR assemblies (Supplementary Table S3). They were present in the eleven CPR phyla contained in the dataset (Figure 1A), and their presence was evenly distributed, regardless of the assembly size (Figure 1B). These VirB4-like proteins were aligned with those from bacterial conjugative plasmids and ICEs of the eight T4SS classes, as well as with VirB4-like proteins representative of archaeal conjugative and DNA-importing systems. Their relationships were assessed using a maximum-likelihood phylogenetic reconstruction.

**Figure 1:**
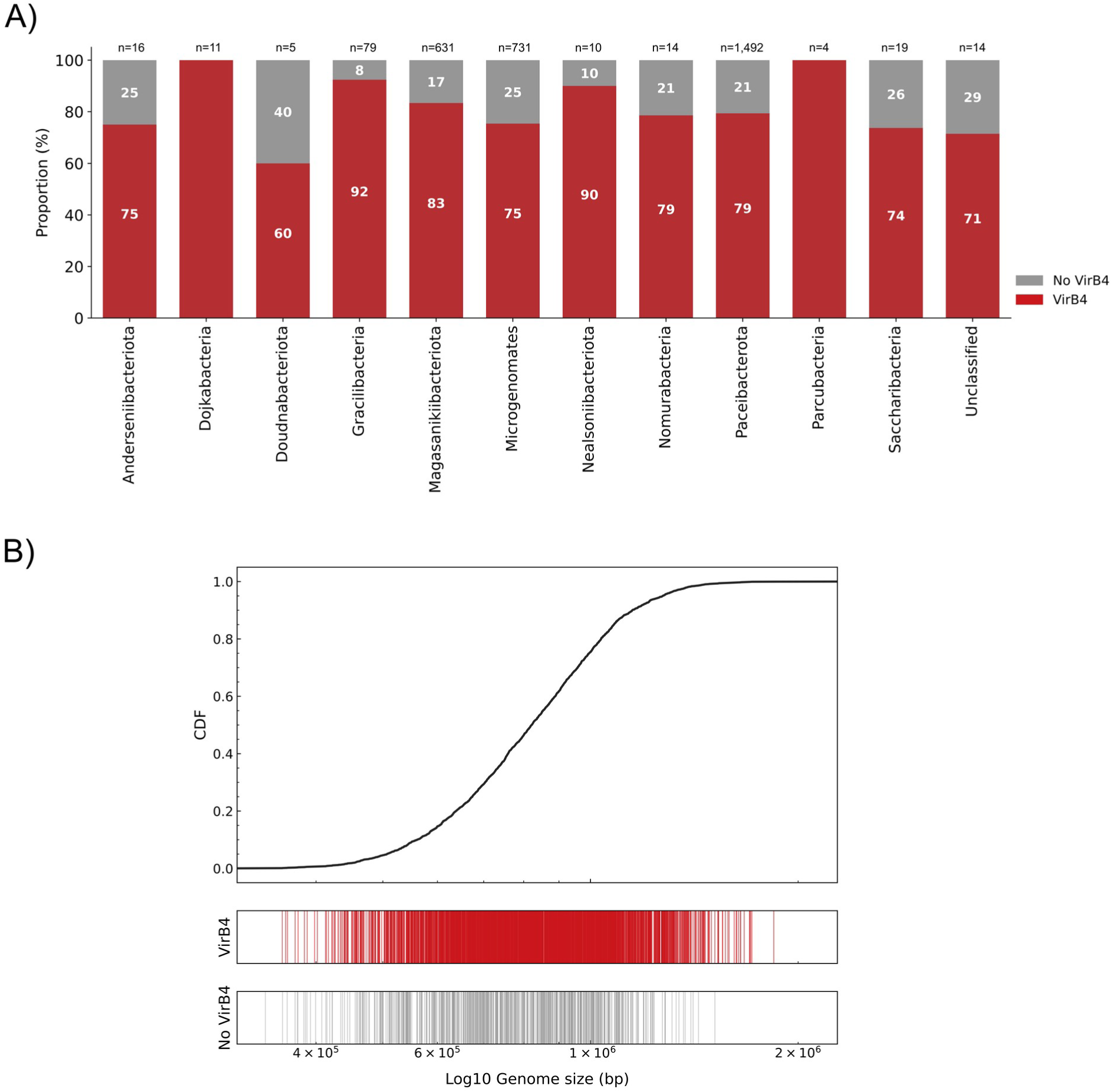
Detection of VirB4 homologs in Patescibacteria. A) VirB4 abundance in the CPR dataset. Stacked bar plot showing the proportion of genomes with (red) and without (grey) VirB4 homologs for each phylum in the dataset. The total number of genomes per phylum (n) is displayed at the top of each bar. **B) Distribution of VirB4 homologs according to genome size.** The 3,026 assemblies were ranked by size (307,478 – 2,280,175 kb), shown on the x-axis. The top panel displays a cumulative distribution function (CDF) plot of genome size, while the lower panels use vertical lines to indicate the presence (red) or absence (blue) of a VirB4 homolog in each corresponding genome.

The resulting phylogeny (Figure 2) showed that a minority of VirB4 homologs present in different phyla of the CPR dataset (49 out of 2,474) clustered together with representative members of the MFF_FATA_ (7 with bacterial VirB4 and 37 with archaeal homologs), MPF_F_ (2), MPF_T_ (2), and MPF_FA_ (1) clades, suggesting that a few CPR bacteria have acquired different T4SS from various sources in independent transfer events. Notably, most VirB4 proteins (2,425 present in 2,407 CPR assemblies) clustered into a monophyletic group without VirB4 homologs from other origins. This CPR clade, supported by 97% SH-aLRT and 98% UF-bootstrap values, is nested within the MPF_FATA_ clade. This phylogenetic positioning, sharing a common ancestor with VirB4 proteins from other monoderms (*Firmicutes*, *Actinobacteria*, *Tenericutes*, and *Archaea*), aligns with the proposed absence of an outer membrane in CPR bacteria (*21*, *58*). It also points to the possibility of a single horizontal transfer event of T4SS from monoderms, likely a basibiont, to a CPR ancestor. However, we cannot exclude the possibility that MPF_FATA_ diversification began before the CPR lineage diverged from other monoderm bacterial phyla.

**Figure 2:**
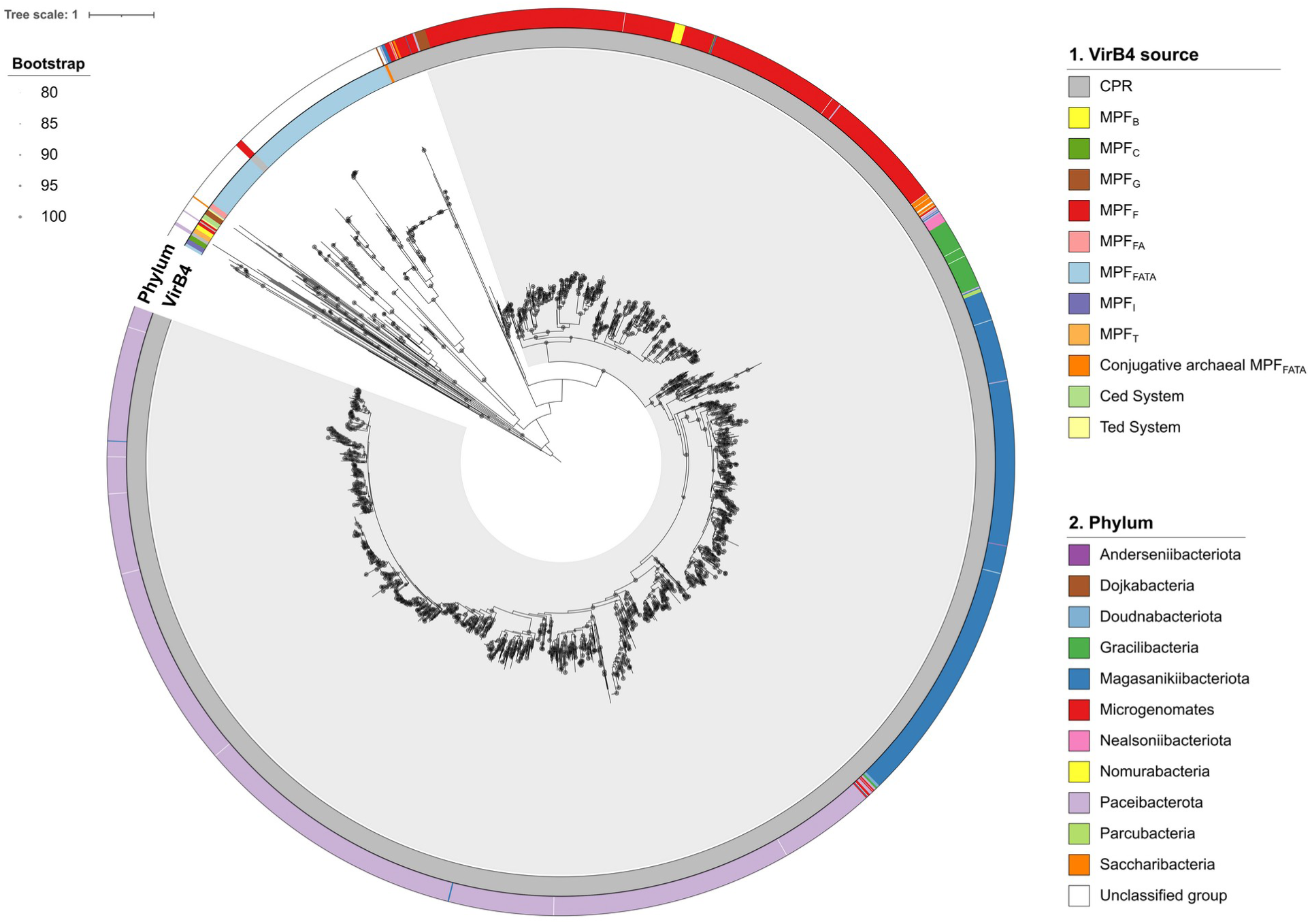
Phylogenetic analysis of VirB4 proteins. Maximum-likelihood tree of the VirB4 homologs of the CPR dataset and members from different T4SS classes rooted at midpoint. Nodes with UFBoot support values >=80% are indicated by a grey circle. Rings from inside to outside indicate: 1) T4SS class, and 2) CPR phylum. The CPR clade is shadowed in grey.

Within the CPR clade, well-defined monophyletic subgroups were observed, each consisting predominantly of VirB4 proteins from a specific CPR phylum. We computed the shortest patristic distance separating each VirB4 protein from the closest homolog in the tree belonging to a different or the same phylum, as a proxy for estimating T4SS exchange between CPR phyla (Figure 3). The cumulative distribution function of patristic distances showed a rapid initial increase for intraphylum comparisons, whereas the increase was slower for VirB4 homologs from different phyla. In fact, 803 proteins have an identical homolog (patristic distance = 0) within the same phylum, while no identical homologs were found in different phyla. Additionally, 99.33% of VirB4 proteins from CPR have their closest homologs within the same phylum, whereas only 0.67% are closest to a homolog from a different phylum. This suggests that, at short evolutionary distances, it is uncommon to find a VirB4 homolog from a different phylum, indicating that T4SS transfer events between CPR phyla are rare.

**Figure 3:**
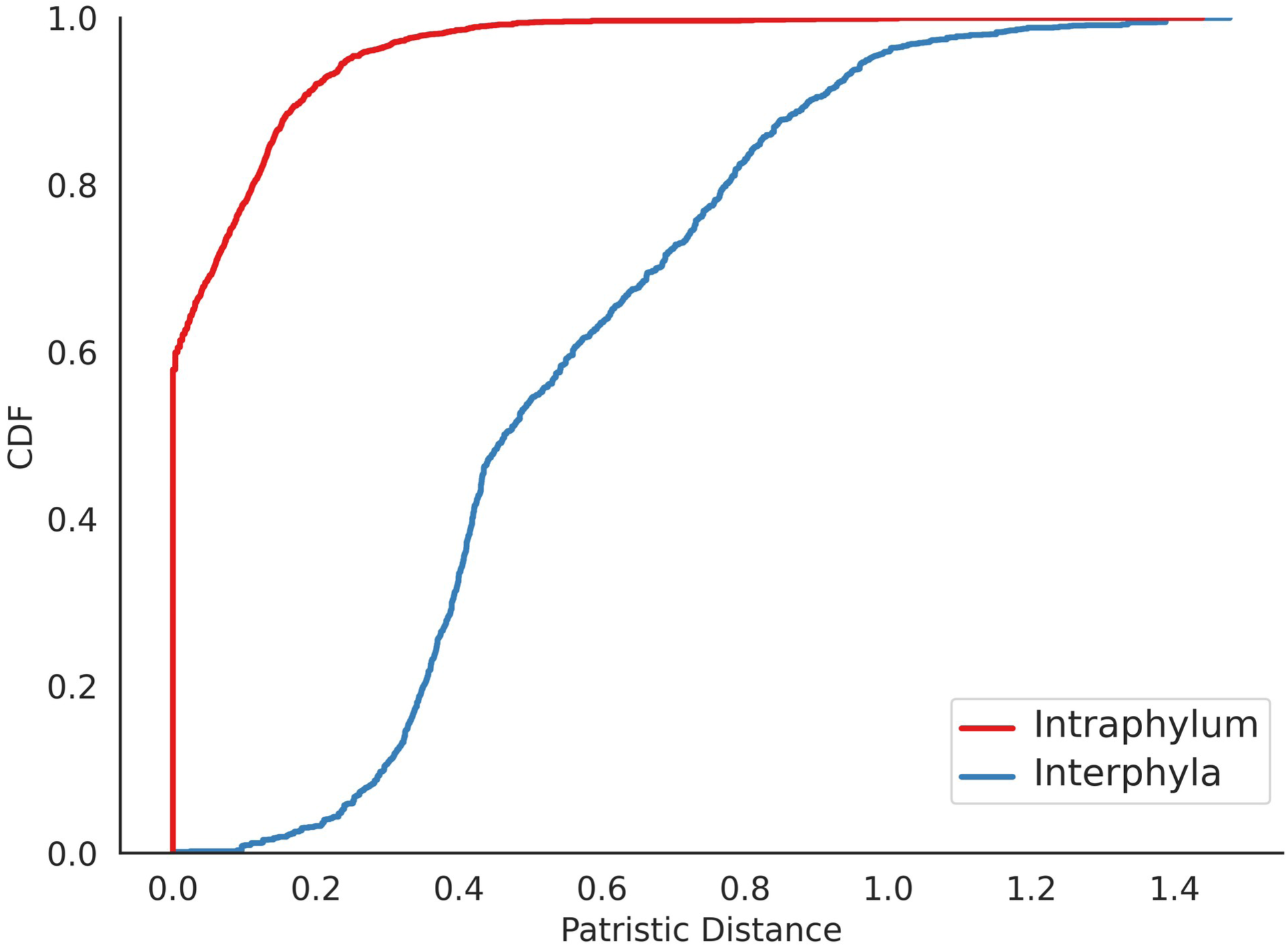
Cumulative distribution function of the minimal patristic distances in the VirB4 tree. The curves show the sum of the lengths of the branches linking two nodes in the VirB4 phylogenetic tree, belonging to the closest homologs within the same phylum (red) or a different phylum (blue).

### T4SS gene repertoire in *Patescibacteria*

Given the phylogenetic proximity of VirB4 proteins in the CPR dataset to those of the MPF_FATA_ type, it is expected that other T4SS components homologous to that system could potentially be found. The MPF_FATA_ class is highly diverse, as are the cell envelopes of the various monoderms included in this class. Moreover, for the MPF_FATA_ class, the exact number of T4SS components, their functions, and their essentiality are still uncertain. A set of HMM profiles covers four MPF_FATA_ subclasses, each based on a different prototype: the plasmids pGO1 (*Staphylococcus aureus*) and pCF10 (*Enterococcus faecalis*), and the ICEs CTn2 (*Clostridium difficile*) and ICESaNEM316 (*Streptococcus agalactiae*) (*40*). We used HMM profile-based searches to detect components of MPF_FATA_ and the other seven T4SS classes. T4SS components are generally encoded near the *virB4* gene (*40*), albeit exceptions have been noted, such as the dispersed T4SS islands throughout *Rickettsia* genomes (*59*), and the coupling protein gene which is close to the relaxase gene in many conjugative systems (*40*). We thus focused primarily on hits associated with VirB4, that is, cases where genes encoding T4SS components are located near the *virB4* gene. T4SS components, primarily associated with MPF_FATA_ subclasses, though not exclusively, were systematically identified in the CPR dataset (Figure 4 and Supplementary Table S3). Notably, many of these components were associated to VirB4. To perform more sensitive searches, CDSs encoded near *virB4* were clustered and an HMM profile (CPR profile) was constructed from the alignment of the members of each cluster. These CPR profiles were compared with the MPF_FATA_ HMM profiles.

**Figure 4:**
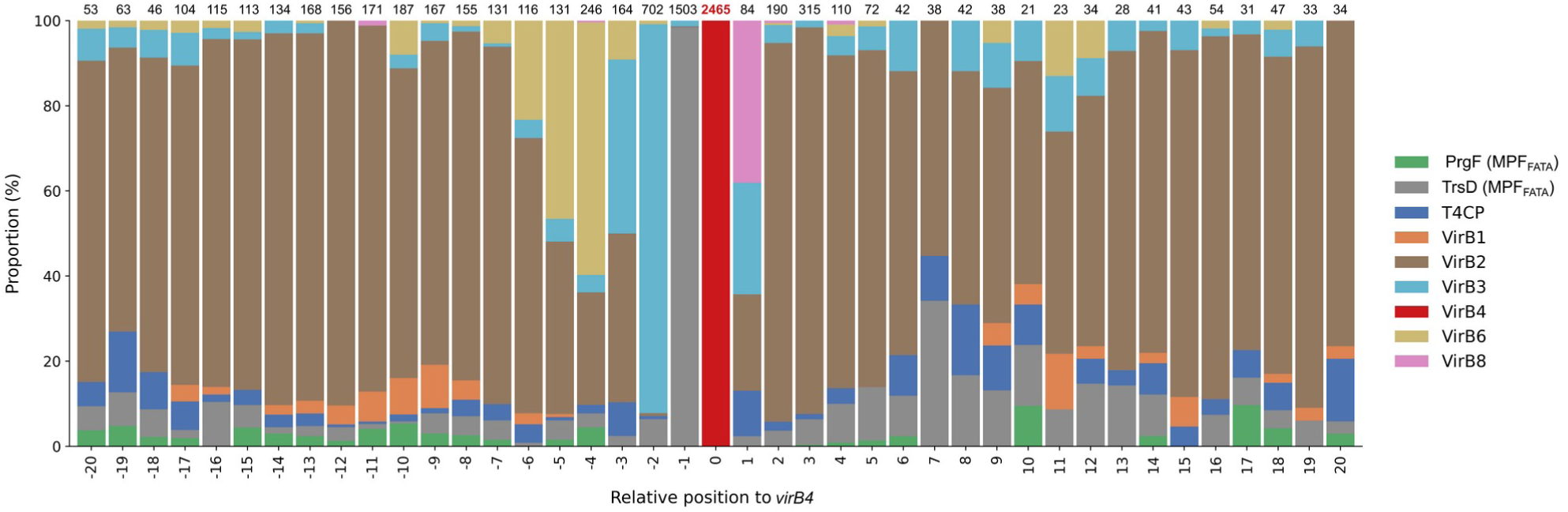
Abundance of T4SS components near *virB4*. Each column represents the abundance of T4SS components at a specific position relative to the *virB4* gene (within a range of -20 to +20 coding sequences). Only *virB4* genes associated to another T4SS component were considered. The total count of T4SS components at each position is shown above each column and represents 100%. The T4SS components are color-coded according to the legend. VirB1 comprises homologs retrieved with HMM profiles MPFFATA TrsG and CD419, and MPFT VirB1; VirB2 those retrieved with HMM profiles MPFB TraE, MPFG Tfc9 and Tfc10, MPFI TraQ and TraR, and MPFT VirB2, and Pfam PF04956.16 and PF18895.3; VirB3 with MPFB TraF, MPFC Alr705, MPFF TraL, MPFFATA PrgI and TrsC, MPFG Tfc11, MPFI TraP and MPFT VirB3; VirB6 with MPF_FATA_ PrgH and MPFT VirB6; and VirB8 with MPFF TraE, MPFFATA PrgL and MPFT VirB8.

All conjugative T4SSs, as well as most T4SSs involved in protein secretion, contain a second ATPase that is ancestrally related to VirB4: a coupling protein (T4CP) (*55*). This protein is located in the inner membrane and acts as a connector between the T4SS channel and the translocated substrates (*2*, *60*). Some exceptions include the *Bordetella pertussis* Ptl system and the *Brucella* sp. VirB system (*56*, *61*), which lack T4CP homologs. Two T4CP families, homologous to VirD4 or TcpA, encompass the diversity of this protein accross the eight T4SS classes (*40*). We detected 5,713 T4CP homologs to VirD4 in 2,918 CPR genomes, with only a minority of them (148) encoded near *virB4* (Supplementary Figure S3). It is not uncommon to find the *t4cp* encoded far from *virB4* (*40*), as observed with the *t4cp* homolog in *Ca. Southlakia epibionticum* strain ML1. However, given that this gene was not found to be essential for its epibiontic growth, while other T4SS genes were (*36*), the possibility of *t4cp* being exapted for other functions cannot be ruled out.

The most abundant T4SS component retrieved from the HMM search analysis was TrsD, a protein that shares remote homology with the N-terminal portion of the VirB4 proteins (*40*). TrsD is present in only one of the four MPF_FATA_ subclasses (prototype pGO1), and no function has been assigned to this protein. A total of 2,272 TrsD homologs were identified across 2,221 CPR assemblies, including 26 detected exclusively through profile-profile alignments (Supplementary Table S3). Most of these homologs (2,209) were found in genomes with VirB4 proteins clustered within the CPR clade. Most of these homologs (1,664) were associated with VirB4 (Figure 4), with the notable exception for TrsD homologs in the *Gracilibacteria* phylum (Supplementary Figure S3). We detected a TrsD homolog associated to VirB4 in *Ca. S. epibionticum* strain *ML1,* which is listed as essential for its survival (*36*).

VirB3, VirB6, and VirB8 are inner membrane proteins that form the cytoplasmic membrane translocon in conjunction with the VirB4 and T4CP ATPases (*62*). VirB3 is an inner membrane protein that interacts with VirB4, anchoring it to the inner membrane (*2*, *56*, *63*). All four MPF_FATA_ prototypes contain VirB3 homologs (*40*). These were recovered in the CPR dataset (1,649 proteins, of which 1,619 in the CPR clade), with a wide variety of HMM profiles from different MPF classes (MPF_FATA_ profiles PrgI (588), and its distant homolog TrsC (97); MPF_F_ profile TraL (142); MPF_T_ profile VirB3 (110); MPF_G_ Tfc11 (53); with MPF_B_ TraF (48) and MPF_C_ Alr1205 (22)) (Supplementary Table S3). HMM-HMM comparisons between CPR and PrgI retrieved 449 additional proteins. VirB3 homologs were mainly associated with VirB4 (880) (Figure 4), except for the homologs from *Microgenomates* and *Gracilibacteria* (respectively 22.6% and 4.7% of them) (Supplementary Figure S4). PrgI has been previously shown to be more abundant in CPR than in non-CPR bacteria (*57*) and was found essential for epibiontic growth in *Ca. S. epibionticum* strain *ML1* (*36*). In some conjugative systems, VirB3 and VirB4 are fused into a single protein (*64*, *65*). However, we found no instances of VirB3-VirB4 fusion proteins in the CPR dataset.

VirB6 is a polytopic integral membrane protein (*66*). Three of the four MPF_FATA_ subclasses contain VirB6 homologs (PrgH/CD415/GBS1362 HMM profiles). However, only 12 out of the 480 homologs detected in the CPR dataset were retrieved with the PrgH HMM profile, while the rest aligned with the VirB6 HMM profile, and most homologs (304) were encoded near *virB4* (Figure 4 and Supplementary Table S3). Two VirB6 homologs were additionally retrieved through CPR-PrgH profile-profile alignments. In *Ca. S. epibionticum* strain *ML1*, the detected *virB6* gene was essential for its growth (*36*). It is noticeable that in well-represented phyla *Microgenomates* and *Gracilibacteria*, VirB6 homologs are respectively scarce or absent (Supplementary Figure S5).

Previous profile-profile comparisons did not identify any MPF_FATA_ HMM profiles with homology to VirB8 (doi: 10.1093/nar/gku194 (*40*)). Nevertheless, structural homologs of VirB8 have been identified in the MPF_FATA_ plasmids pIP501 (TraM (*67*) and TraH (*68*)) and pCF10 (PrgL (*69*) and PrgD (*67*)). We detected a few instances of homology to VirB8 HMM profiles from different T4SS classes (MPF_T_ VirB8 (20 proteins), MPF_F_ TraE (8), and MPF_FATA_ PrgL (1), most associated with VirB4 (Supplementary Table S3 and Supplementary Figure S6). More sensitive searches using CPR-PrgL profile comparisons detected 22 additional homologs. Considering the limited number of VirB8 homologs identified, nearly all of which were found in a single phylum, *Magasanikiibacteriota*, it is likely that this protein has undergone substantial diversification within CPR bacteria.

VirB1 is another protein commonly found in T4SSs. This protein is not part of the macromolecular complex but aids in the assembly of the T4SS channel by locally degrading the peptidoglycan (*70*). In some monoderms, *virB1* genes are significantly larger than those in diderms (*62*), as they contain two domains: an N-terminal soluble lytic transglycosylase domain (SLT) and a C-terminal cysteine-, histidine-dependent amidohydrolase/peptidase (CHAP) domain. Both domains are involved in degrading peptidoglycan (*71–75*). VirB1 homologs were retrieved with the HMM profiles MPF_T_ VirB1 (15), MPF_FATA_ TrsG (8) and CD419 (74); and through HMM-HMM comparison between CPR HMM and CD419 (72) and TrsG (11) (Supplementary Table S3 and Supplementary Figure S7). In most phyla, except for *Paceibacterota*, VirB1 homologs were not highly associated with VirB4. We cannot rule out the possibility that other glycosidases might fulfill the role of VirB1 in CPR. This appears to be the case in *Ca. Southlakia epibionticum* strain ML1, where peptidase and lysozyme homologs encoded immediately downstream of *virB4* have been found essential for its epibiotic growth (*36*).

In monoderms, T4SS classes lack homologs for the outer membrane core complex proteins (VirB7, VirB9, and VirB10) found in diderm T4SSs (*1*). Yet PrgC, which is exclusive to one of the four MPF_FATA_ subclasses, shows distant homology to the N-terminal region of VirB9 (*40*), which is known to interact with the inner membrane (*76*). We detected 56 instances of VirB9 homologs, of which only seven encoded near *virB4* (Supplementary Table S3 and Supplementary Figure S8). In the CPR dataset, no other T4SS components of the outer membrane were associated with VirB4, aside from two VirB10 homologs in assemblies whose VirB4 proteins were located within the MPF_T_ or MPF_F_ clade, respectively, and a third homolog present in the CPR clade (Supplementary Table S3). Nevertheless, T4SS classes in monoderms (MPF_FATA_ and MPF_FA_) encode, near the *virB4* gene, other components specific to each class that are absent in other T4SS types and have no known function. In CPR, a few were detected, including PrgF (627 proteins), TrsJ (47), Gbs1347 (20) and Gbs1350 (37) from MPF_FATA_ and Orf17 (109) from MPF_FA_ (Supplementary Table S3).

The T4SS classes present in Gram-positive bacteria lack T4SS major pilin genes (*virB2* homologs), relying instead on surface-exposed adhesins or outer membrane proteins to establish and stabilize donor-recipient contacts (*14*, *77*, *78*). The type IV pili (T4P), although not a component of the T4SS, collaborates with it by facilitating mating-pair formation rather than directly transporting DNA. T4P encoded in certain conjugative plasmids, such as the IncI plasmid R64, is crucial for conjugative transfer, though only in liquid mating conditions (*79*). Similarly, in Archaea it has been shown to be essential for DNA import by the Ced system (*15*, *17*). In CPR, T4P was also found to be essential for enabling adhesion to the host during episymbiosis (*30*). No adhesins, such as the MPF_FATA_ PrgB, were detected in the CPR dataset. Considering that a VirB2 homolog was identified in the parasitic *Saccharibacteria Ca. Nanosynbacter lyticus* strain TM7x (*35*), we conducted a search for VirB2 homologs using various HMM profiles, taking into account the small size of this protein. Using pilin HMM profiles from different T4SS classes of diderms, we detected VirB2-like pilin proteins in 1,948 CPR assemblies that also contained VirB4, with the pilin gene located near *virB4* in 1,135 of these assemblies (Figure 4, Supplementary Table S3 and Supplementary Figure S9). Notably, 441 VirB2 arrays containing 2–11 pilin genes were detected in 383 assemblies. These genes are arranged either in tandem or separated by a non-pilin gene. They include the previously reported essential pilin array in *Ca. S. epibionticum* strain ML1 (*36*) and what was referred to as Sec secreted array found in *Ca. N. lyticus* strain TM7x (*35*). These VirB2 arrays were especially abundant in *Microgenomates* and *Saccharibacteria* (Supplementary Figure S9). The VirB2 proteins within each array showed high variability, with an average amino acid identity of just 25% and a maximum identity of 54%. Examples of tandem amplification and variation of pilin genes are found in virulence-associated T4SS of intracellular bacteria, such as different species of *Bartonella* (*80–82*) and *Anaplasma phagocytophilum* (*65*). In such cases, the presence of multiple VirB2 paralogs is thought to support a broader immune evasion strategy via antigenic variation or may enhance interactions with a range of host cell surface structures (*81*). For CPR bacteria, this versatility might aid in adapting to different host strains.

Conjugation-related T4SSs are associated with a relaxase (*56*), and nine relaxase MOB classes are distinguished (*42*). We only detected 109 relaxase homologs in 103 CPR genomes, distributed across eight relaxase classes: MOB_C_ (58), MOB_M_ (36), MOB_P_ (8), MOB_V_ (2), MOB_Q_ (2), MOB_T_ (1), MOB_F_ (1), and MOB_H_ (1). In most cases (105 out of 109), the relaxase was identified in a *virB4*-encoding CPR genome and in only 15 of them were the relaxase and *virB4* genes close (Supplementary Table S3 and Supplementary Figure S10). These data suggest that CPR bacteria have acquired mobile genetic elements from different sources through conjugation, and thus only a minority of them have conjugative potential. Therefore, the T4SS identified in the CPR clade appears to be functionally specialized for roles other than conjugation, unless very different relaxases are yet to be found in these bacteria.

### The chromosomal context of T4SS in *Patescibacteria*

The presence of T4SS components near *virB4* were also identified by homology searches against protein family profiles of other databases (Figure 5A, 5B). Besides, we identified several other gene families that are abundant in the vicinity of the *virB4* gene (Supplementary Table S4). For each phylum, the distribution of the six most abundant protein families from each HMM database is shown in Figure 5, and the synteny of the *virB4*-containing region in representative complete genomes of the four most abundant phyla is illustrated in Figure 6. These findings suggest that the genomic location of T4SS varies across different CPR phyla. Among most phyla, including the most represented *Paceibacterota* and *Microgenomates*, genes encoding proteins related to DNA replication, such as the beta subunit of DNA polymerase III DnaN and the chromosomal replication initiator protein DnaA are abundantly represented near *virB4*. Ribosomal proteins, such as the ribonuclease P protein component RnpA, the ribosomal RNA small subunit methyltransferase A KsgA, and the 50S ribosomal protein L34 RpmH are also abundant in the *virB4* vicinity. In many bacterial species, the replication genes, *dnaA* and *dnaN* (*84*), along with ribosomal protein genes (*85*), are typically located near the origin of replication. Bacteria that undergo overlapping rounds of replication tend to have a higher abundance of genes situated close to the origin than those near the replication terminus (*86*). This phenomenon, known as the replication-associated gene dosage effect, results in increased expression of genes located near the origin of replication (*87–91*). Therefore, it is plausible to suggest that the proximity of T4SS genes to the putative origin of replication, at least in some CPR phyla, may play a role in enhancing the expression of these T4SS components.

**Figure 5:**
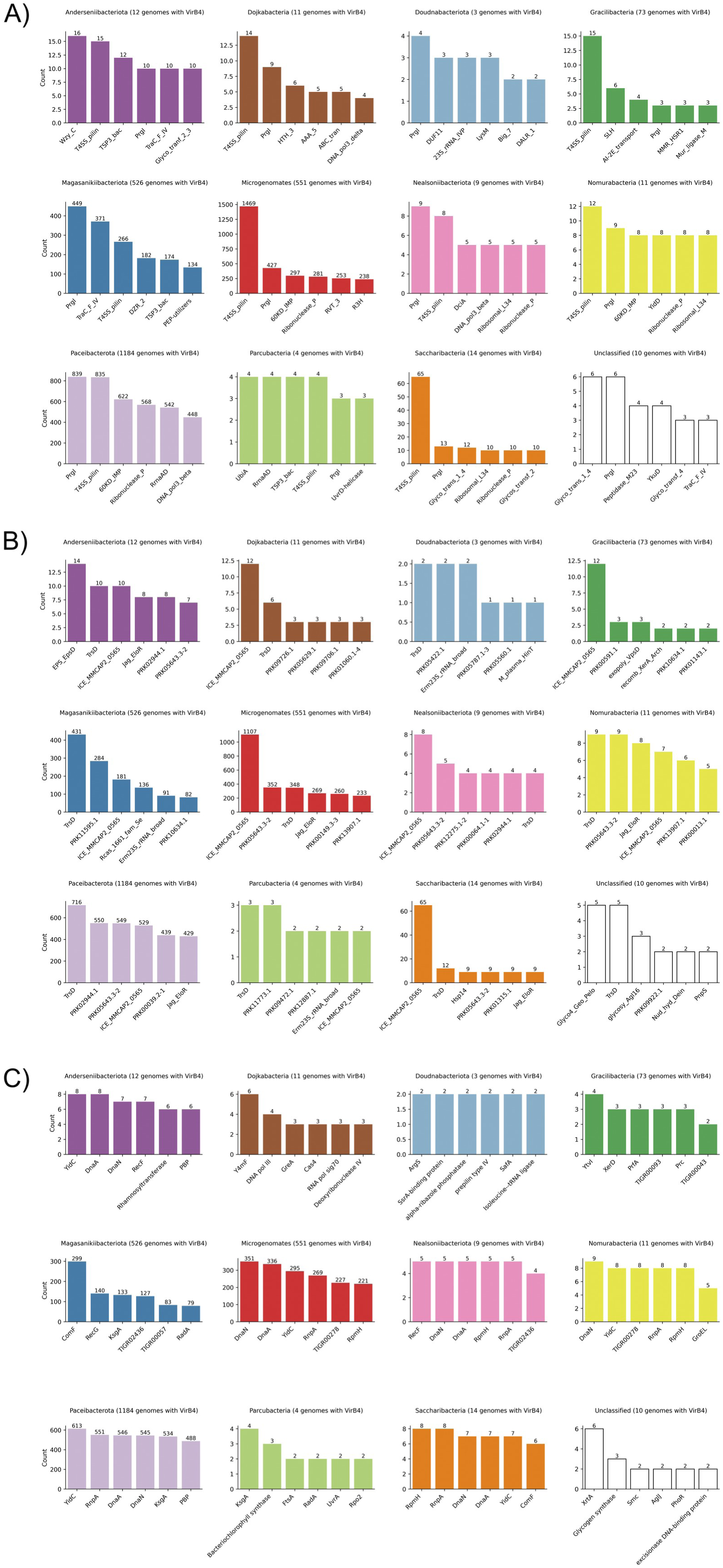
Proximity of the *virB4* gene to other chromosomal functions. For each phylum, the six most prevalent A) Pfam, B) NCBIFAM, and C) TIGRFAM families identified near *virB4* (within a range of -20 to +20 coding sequences) are shown. The count of detected members for each protein family is displayed above the respective bar, while the total number of genomes analyzed per phylum is also indicated.

**Figure 6:**
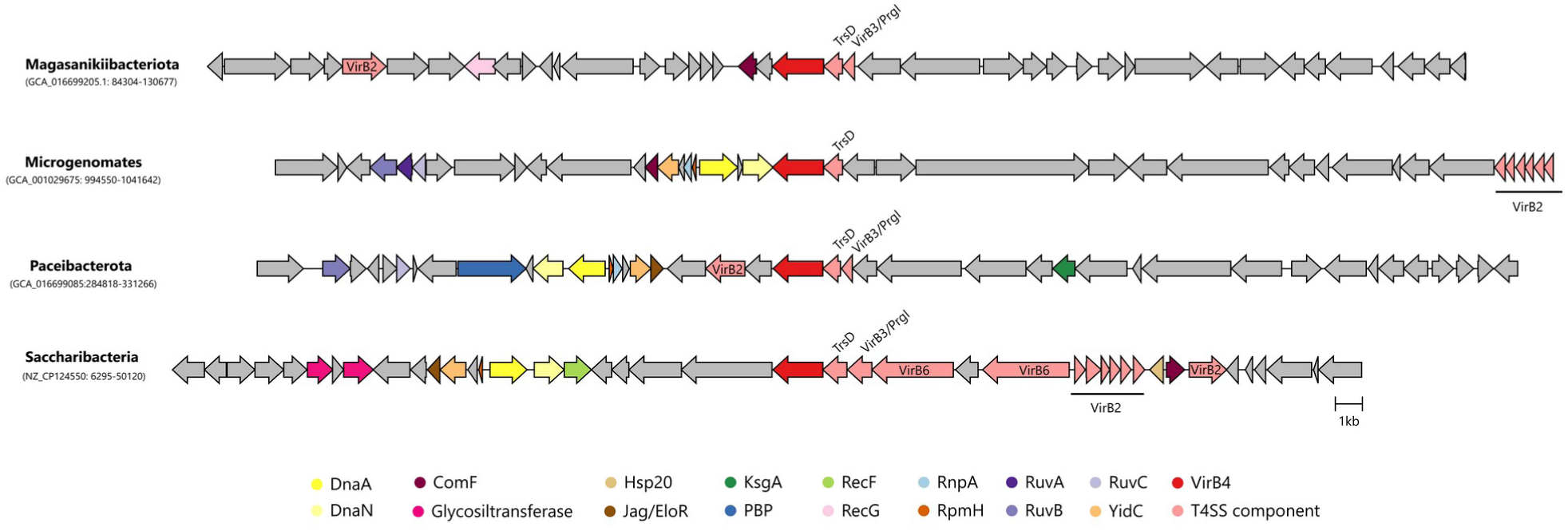
Genomic context of T4SS in representative genomes. For the four most represented CPR phyla in which a T4SS was identified, the synteny of the *virB4* gene neighborhood is illustrated using a representative from complete CPR genomes. The genetic organization encompasses 20 CDSs upstream and downstream of the *virB4* gene (colored in red and located at the center). T4SS genes other than *virB4* are colored in light red, and non-T4SS genes are depicted in different colors according to the legend. Genes for which no homology to NCBIFAM, Pfam-A, or TIGRFAM profiles were found are colored in grey.

On the other hand, in the phylum *Magasanikiibacteriota*, which is also well-represented, genomic loci containing T4SS genes also commonly include genes related to natural competence (*comF*) and DNA recombination (*recG* and *radA*). The co-localization of these genes suggests a functional coordination of their activities.

## Concluding remarks

In this study, we identified the presence of a T4SS in CPR bacteria across several CPR phyla, including *Microgenomates, Paceibacterota, Gracilibacteria, Anderseniibacteriota, Magasanikiibacteriota, Dojkabacteria, Nealsoniibacteriota, Nomurabacteria, Parcubacteria, <u>Doudnaibacteriota</u>,* and *Saccharibacteria*. Homologs of VirB4, TrsD, PrgI/VirB3, VirB6, and VirB2-like proteins were found here to be widely distributed in *Patescibacteria*. Considering the reduced genomes of CPR bacteria, it is notable that they have not only retained the T4SS but often located it at a super-expressed location near the origin of replication. The single experimental analysis of a T4SS carried out in CPR showed that genes encoding these components were essential for the epibiotic host-dependent lifestyle (*36*).

CPR bacteria, lacking many biosynthetic pathways, presumably grow using molecules derived from active hosts. Notably, although genes encoding homologs of the natural competence *comEC* system mediate DNA uptake from the environment, they were not essential for the epibiotic growth of *Ca. S. epibionticum* strain ML1 (*36*), suggesting that CPR bacteria likely depend on alternative systems to acquire nucleotides needed for growth.

T4SSs act as versatile nanomachines, adapted to transfer large macromolecules across multiple cell membranes. The structural adaptability of T4SS has given rise to a broad range of system variations across bacterial lineages, with T4SS having been co-opted repeatedly throughout evolutionary history to enable the import or export of various substrates (*55*), including the acquisition of DNA from the environment or other cells. It is therefore probable that CPR bacteria have repurposed the monoderm conjugative system for functions other than conjugation. The wide distribution and abundance of T4SS in CPR phyla, along with its probable high expression due to the proximity of T4SS genes to the origin of replication, suggest that T4SS in CPR may function as a mechanism for importing DNA or other macromolecules from their hosts, enabling them to exist as obligate episymbionts on other microbes. Future research should explore this hypothesis.

