## Supplementary Figures for "“The Type IV Secretion System of *Patescibacteria* is homologous to the bacterial monoderm conjugation machinery”"

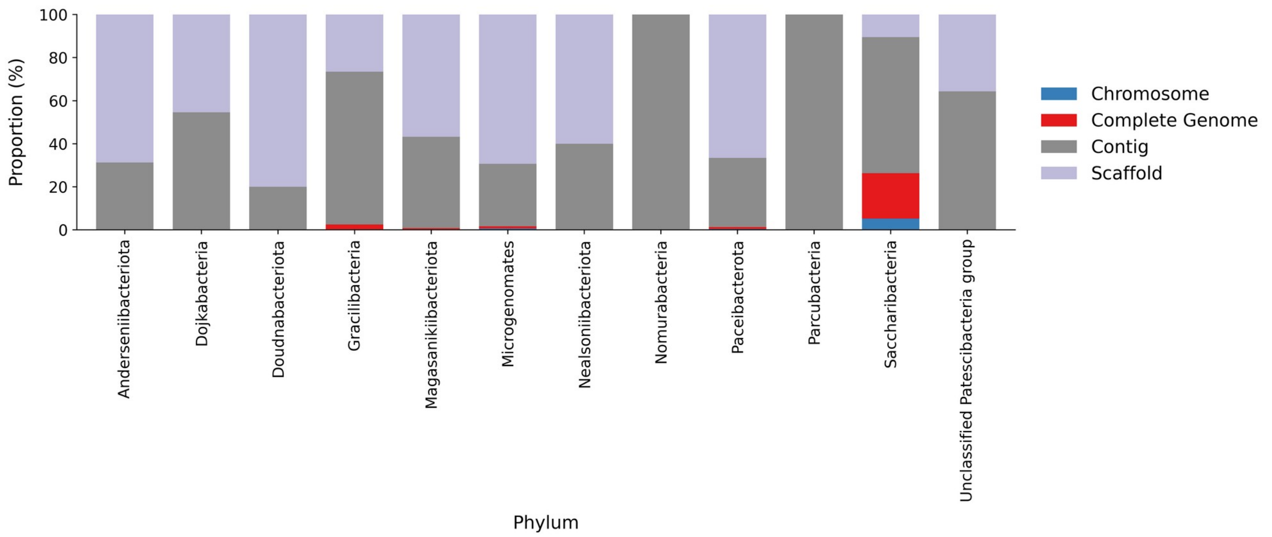


**Supplementray Figure S1: Distribution of the *Pastescibacteria* dataset according to the assembly level.** The proportion of genomes at different assembly levels for each *Patescibacteria* phylum is displayed as stacked bars, with colors corresponding to the legend.

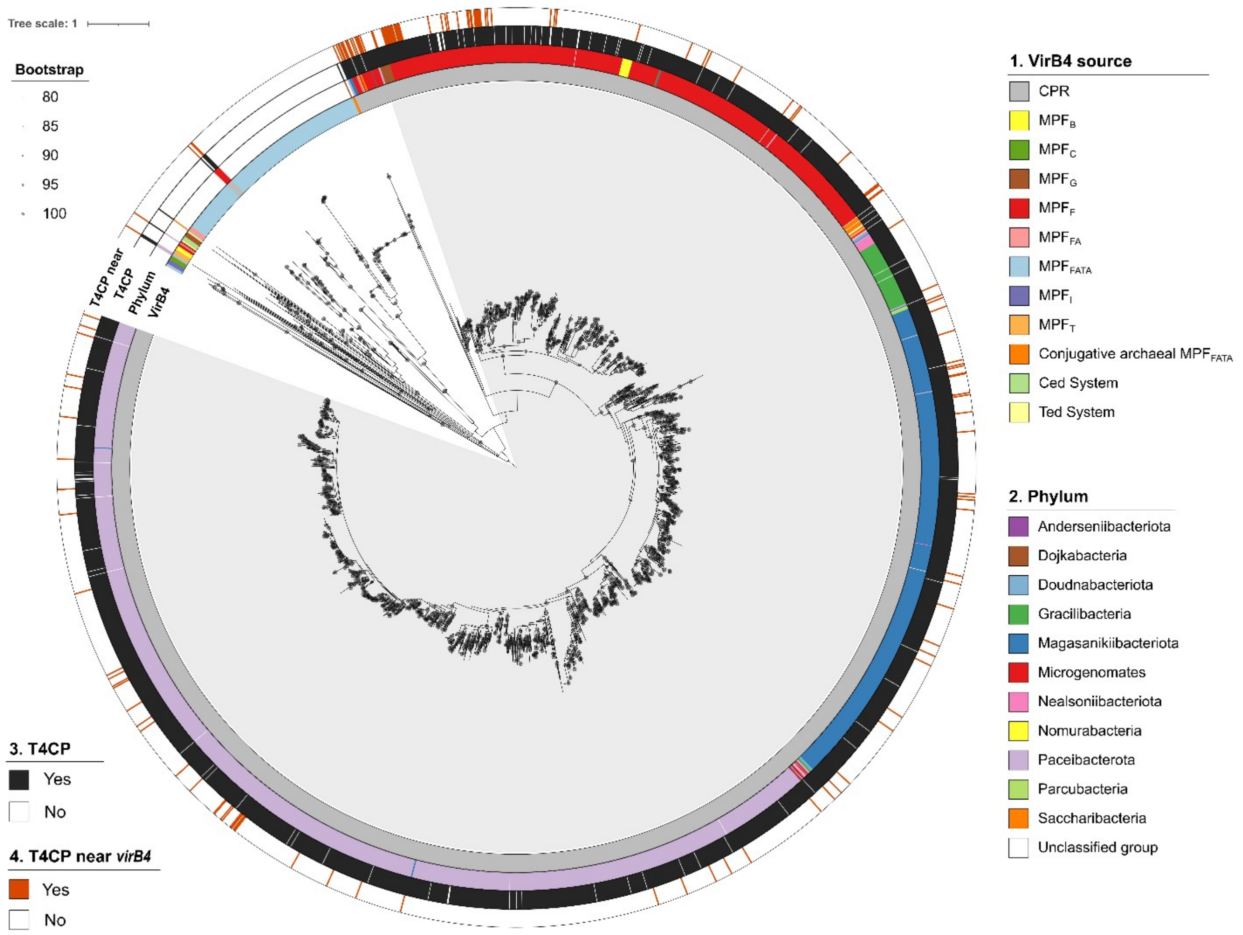


**Supplementary Figure S2: Maximum-likelihood tree of VirB4 proteins.** The tree, along with rings 1 and 2, is as shown in Figure 2. Ring 3 displays the presence of T4CP homologs, while ring 4 indicates those encoded near *virB4*.

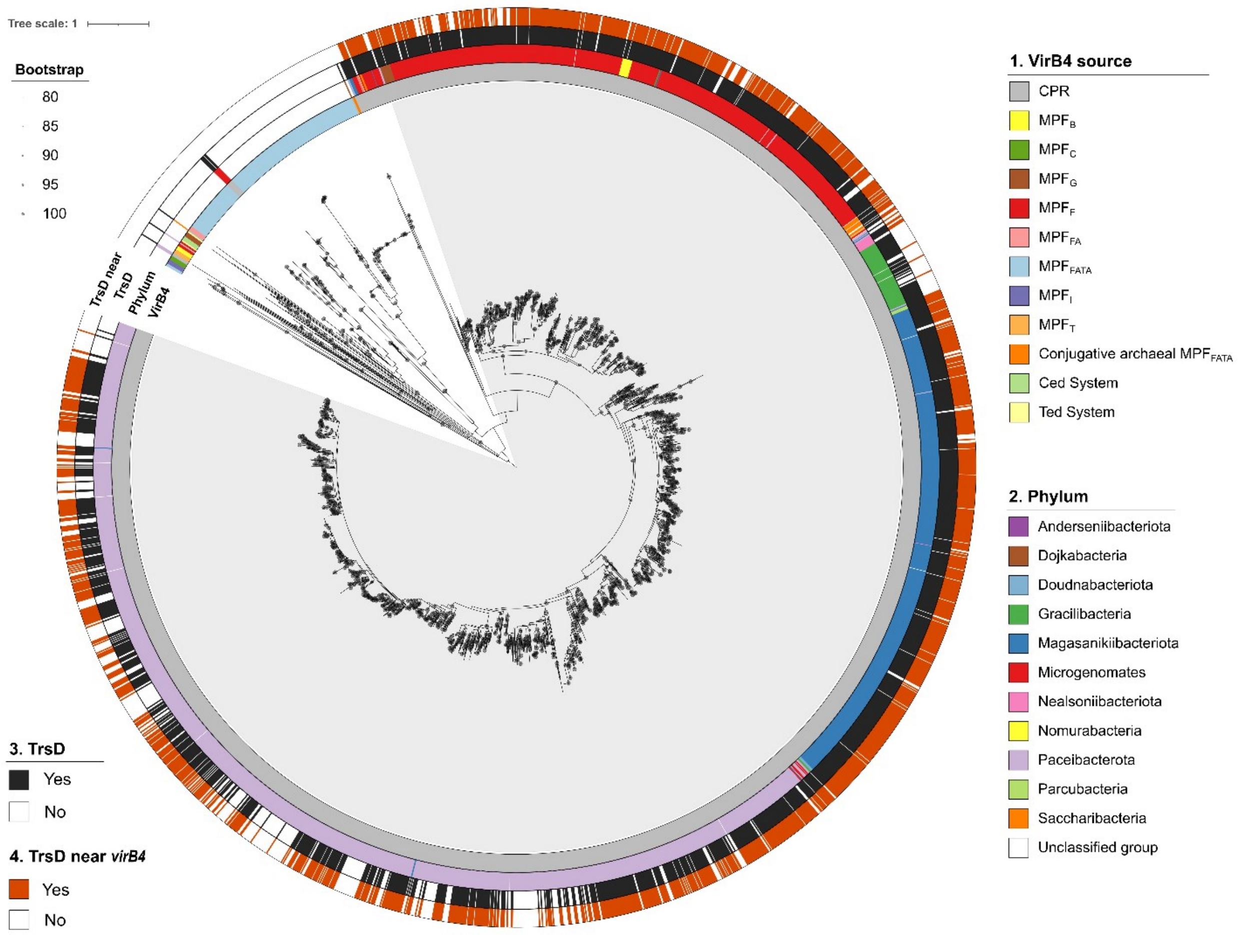


**Supplementary Figure S3: Maximum-likelihood tree of VirB4 proteins.** The tree, along with rings 1 and 2, is as shown in Figure 2. Ring 3 displays the presence of TrsD homologs, while ring 4 indicates those encoded near *virB4*.

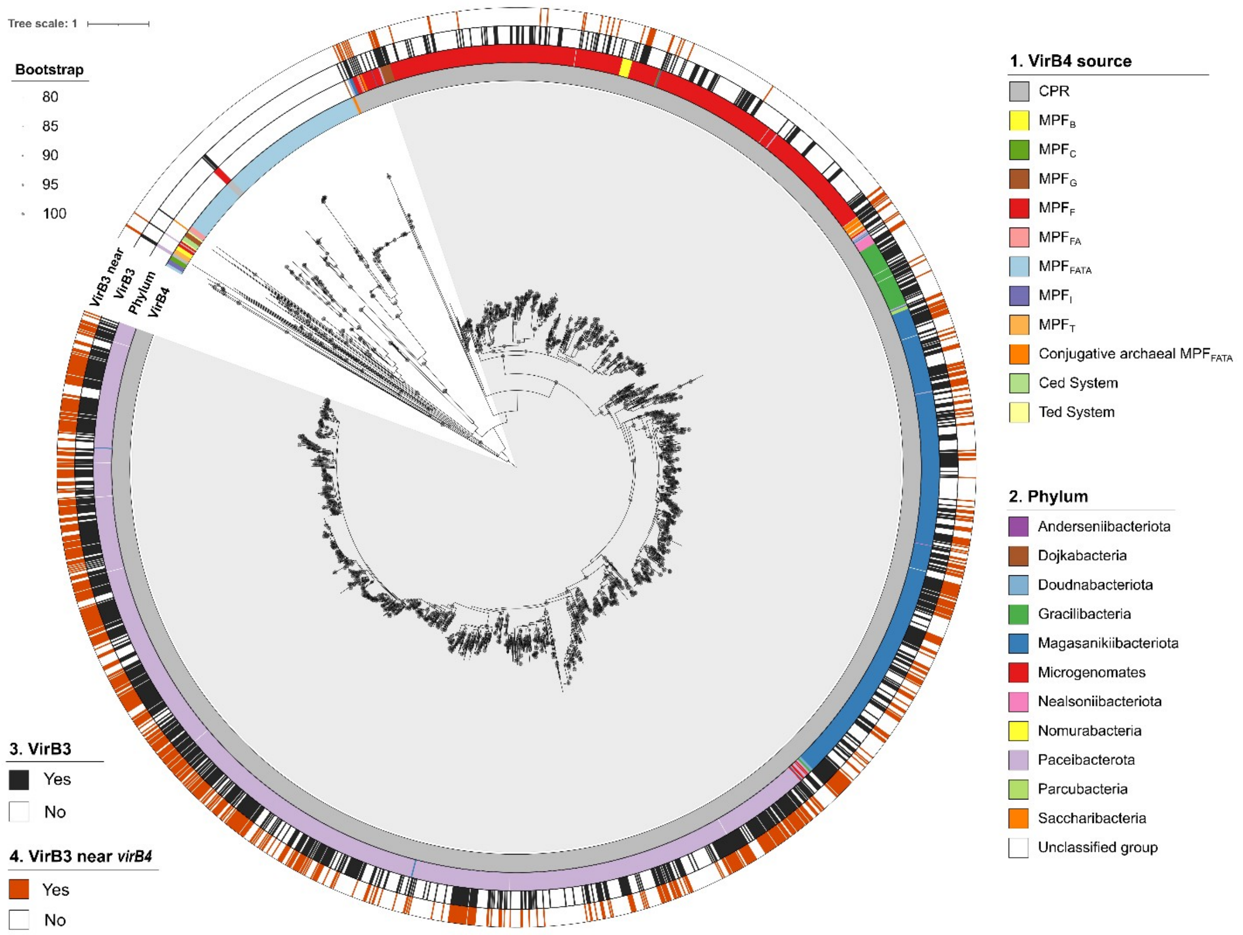


**Supplementary Figure S4: Maximum-likelihood tree of VirB4 proteins.** The tree, along with rings 1 and 2, is as shown in Figure 2. Ring 3 displays the presence of VirB3 homologs, while ring 4 indicates those encoded near *virB4*.

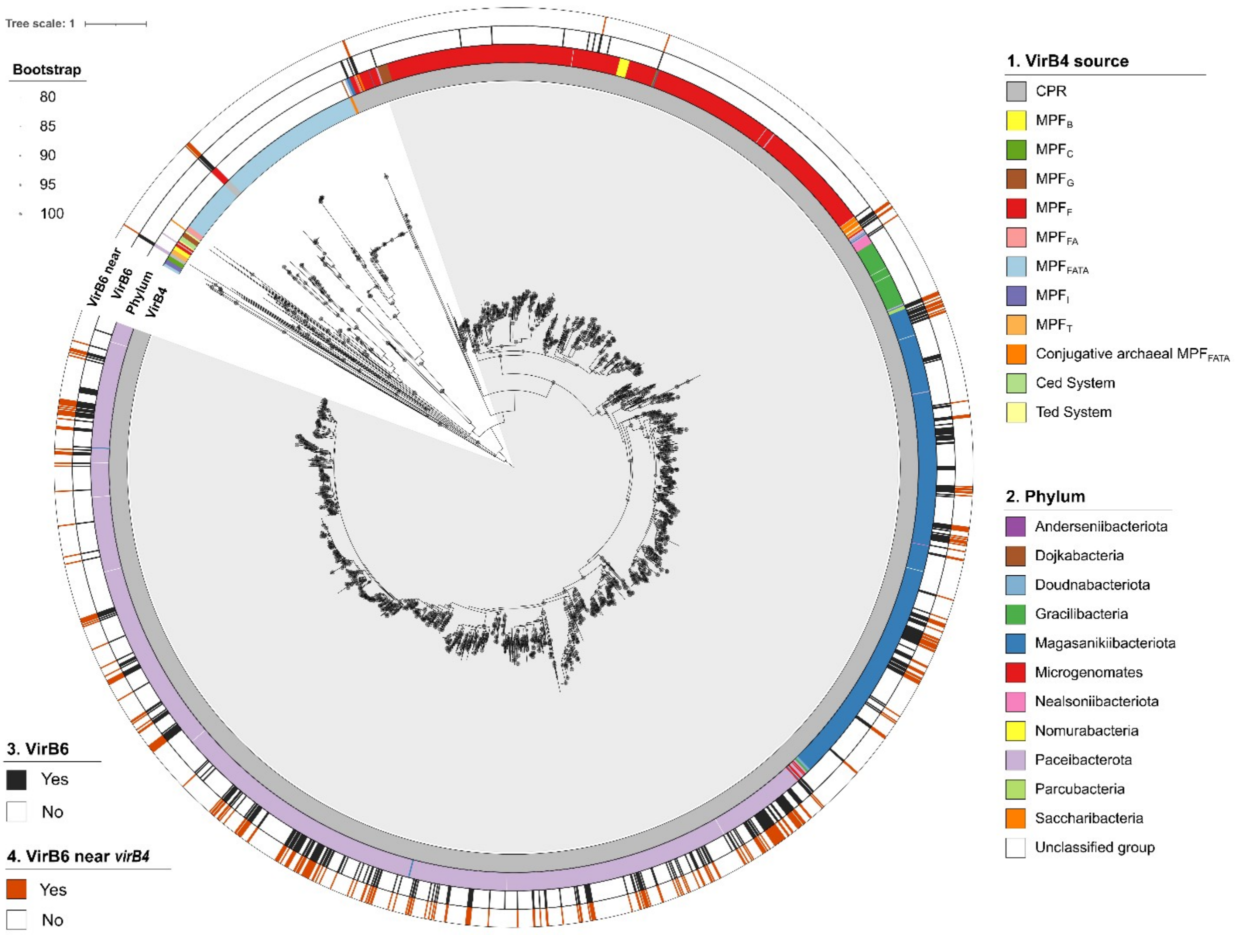


**Supplementary Figure S5: Maximum-likelihood tree of VirB4 proteins.** The tree, along with rings 1 and 2, is as shown in Figure 2. Ring 3 displays the presence of VirB6 homologs, while ring 4 indicates those encoded near *virB4*.

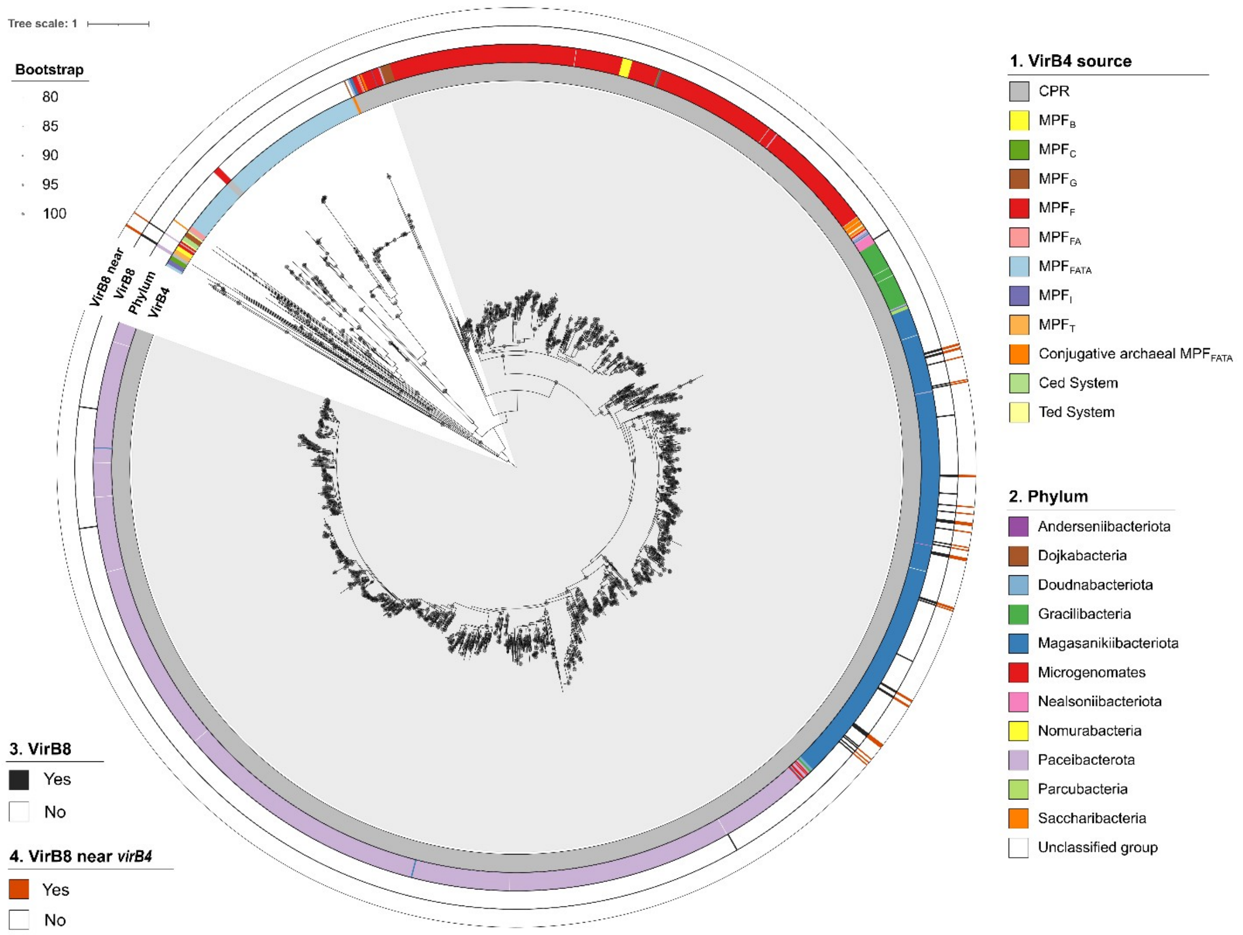


**Supplementary Figure S6: Maximum-likelihood tree of VirB4 proteins.** The tree, along with rings 1 and 2, is as shown in Figure 2. Ring 3 displays the presence of VirB8 homologs, while ring 4 indicates those encoded near *virB4*.

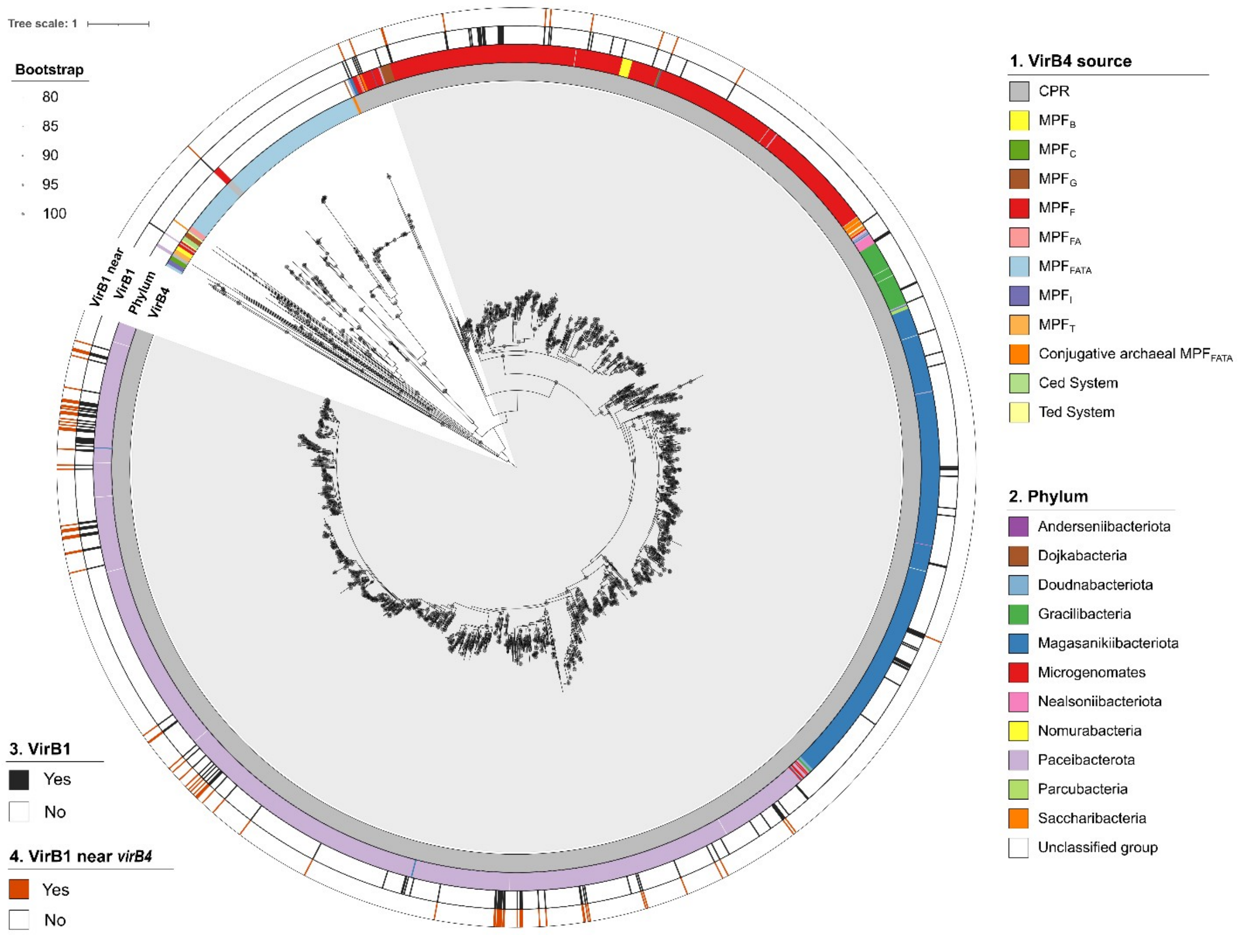


**Supplementary Figure S7: Maximum-likelihood tree of VirB4 proteins.** The tree, along with rings 1 and 2, is as shown in Figure 2. Ring 3 displays the presence of VirB1 homologs, while ring 4 indicates those encoded near *virB4*.

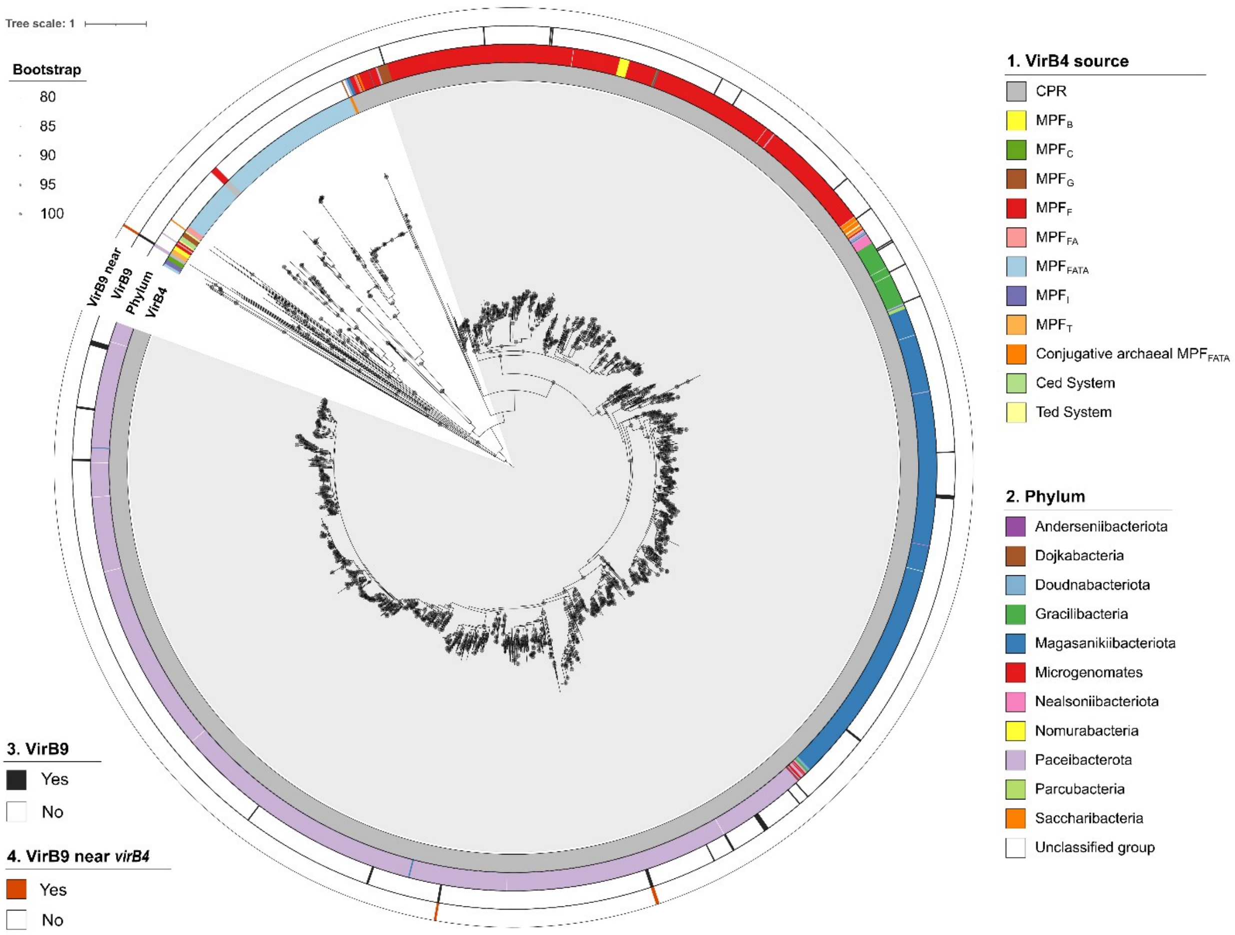


**Supplementary Figure S8: Maximum-likelihood tree of VirB4 proteins.** The tree, along with rings 1 and 2, is as shown in Figure 2. Ring 3 displays the presence of VirB9 homologs, while ring 4 indicates those encoded near *virB4*.

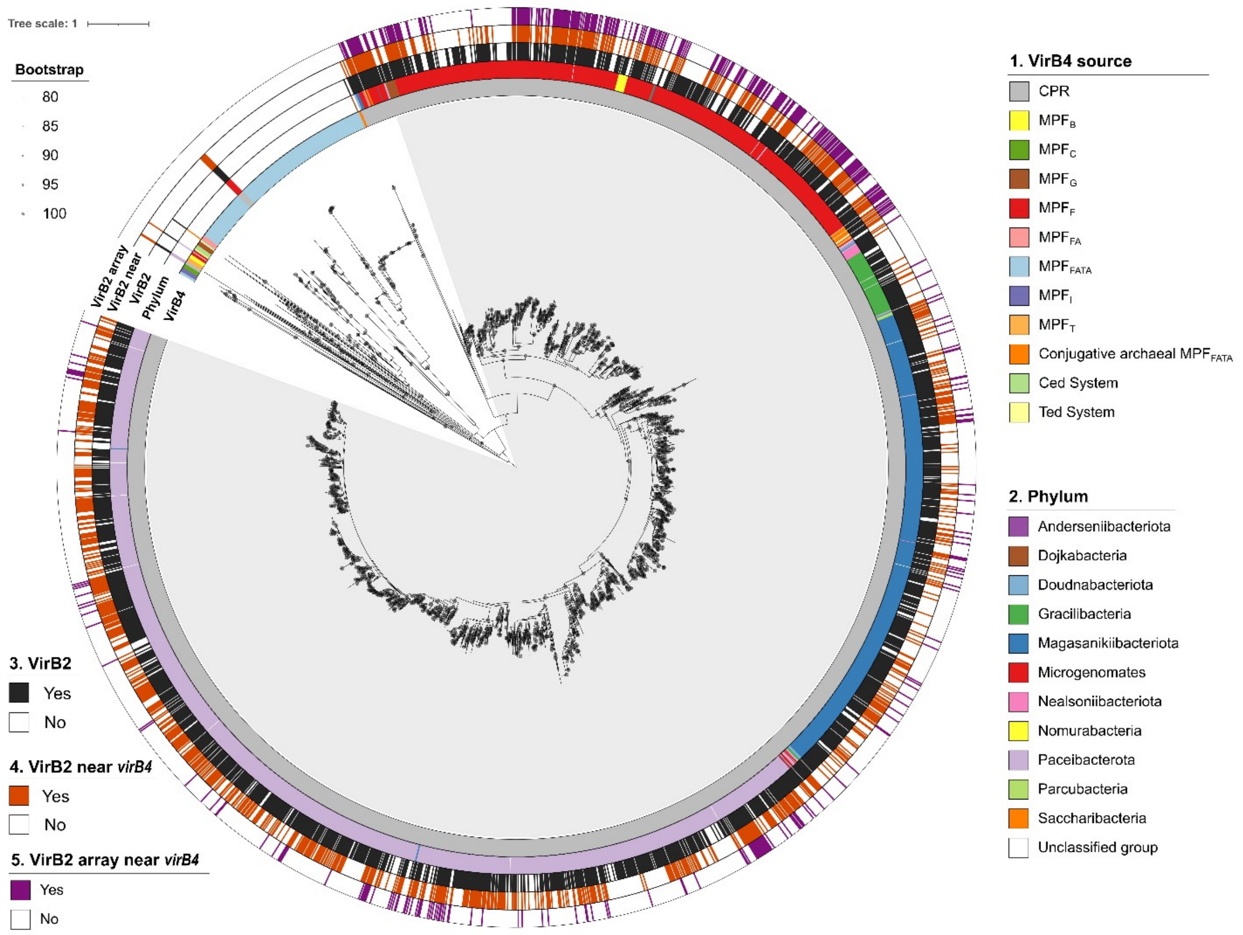


**Supplementary Figure S9: Maximum-likelihood tree of VirB4 proteins.** The tree, along with rings 1 and 2, is as shown in Figure 2. Ring 3 displays the presence of VirB2 homologs, ring 4 indicates those encoded near *virB4* and ring 5 shows the presence of a VirB2 array near *virB4*.

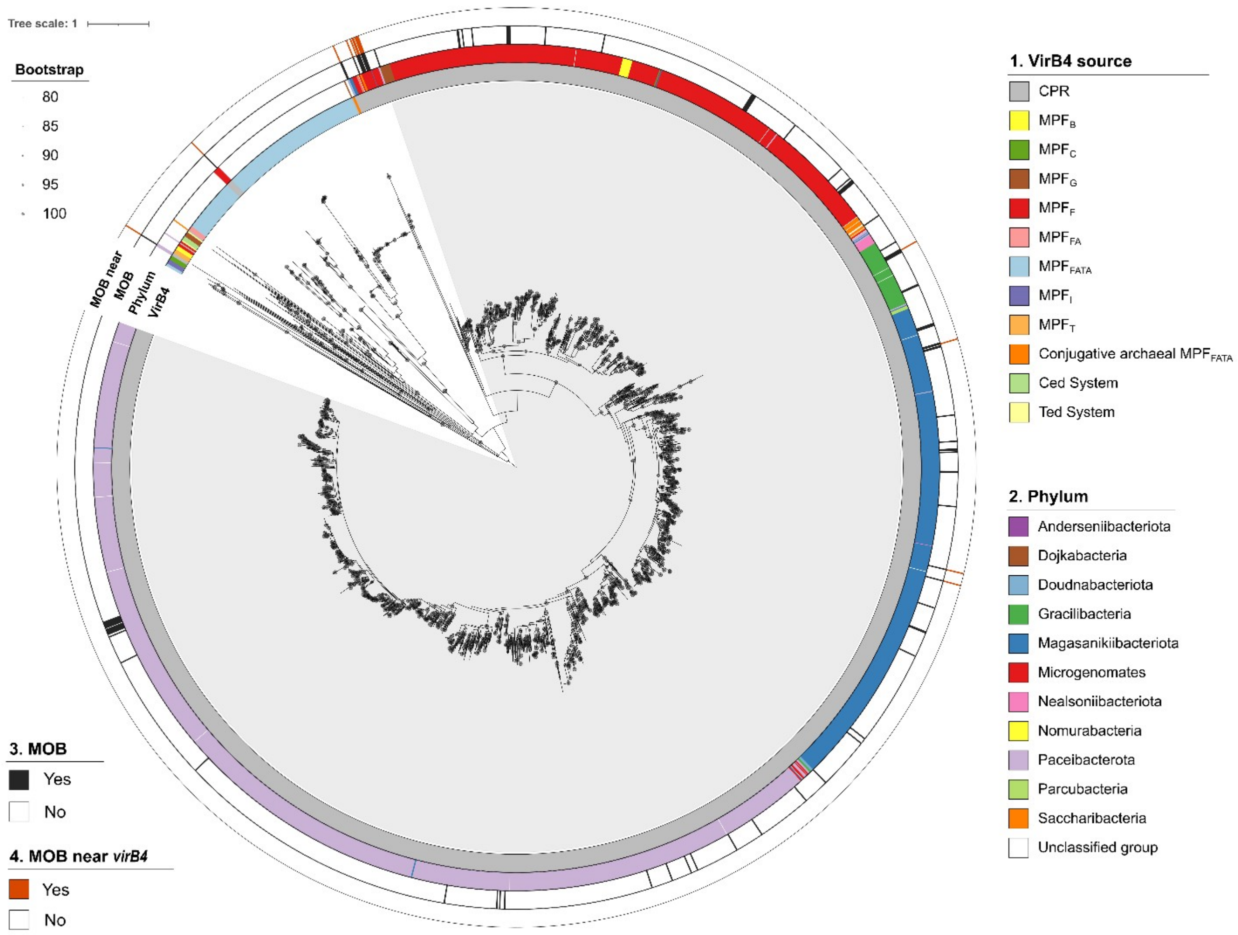


**Supplementary Figure S10: Maximum-likelihood tree of VirB4 proteins.** The tree, along with rings 1 and 2, is as shown in Figure 2. Ring 3 displays the presence of MOB relaxase homologs, while ring 4 indicates those encoded near *virB4*.
